# Specific binding modes of SIRT6 C-terminal domain to the nucleosome core particle influence DNA unwrapping and H3K27 accessibility

**DOI:** 10.1101/2025.04.02.646764

**Authors:** Yuya Qiu, Gabor Papai, Adam Ben Shem, Emmanuelle Bignon

**Author notes:** Corresponding Author Emmanuelle Bignon.

## Abstract

Sirtuins are a class of NAD-dependent histone deacetylases that regulate important biological pathways in prokaryotes and eukaryotes. This enzyme family comprises seven members, named SIRT1 to SIRT7. Among them, Sirtuin 6 (SIRT6) is a human sirtuin that deacetylates histones and plays a key role in DNA repair, telomere maintenance, carbohydrate and lipid metabolism, and lifespan. SIRT6’s structure consists of a zinc finger domain, a Rossmann fold domain containing the NAD+ binding site, and disordered N-terminal and C-terminal (CTD) extensions. The specific role of the CTD on SIRT6 interaction with nucleosomes for histone deacetylation remains unclear. Here, we resort to extended molecular dynamics simulations to uncover the dynamical behavior of the full-length SIRT6 bound to a nucleosome core particle. Our simulations reveal that the CTD preferentially interacts with DNA at the entry/exit near the enzyme’s docking site, exhibiting a variety of different binding modes. In specific cases, the CTD participates to the promotion of DNA unwrapping and promotes H3K27 accessibility to SIRT6’s active site, suggesting a pivotal role of this domain for H3K27 deacetylation. This work provides new structural insights into the binding process of the full-length SIRT6 to a nucleosome core particle, shedding light on the conformational behavior and functional role of its CTD. It constitutes an important step towards the understanding of SIRT6 deacetylation mechanisms and specificity.

## INTRODUCTION

Sirtuins constitute a multifunctional family of proteins found in eukaryotes and prokaryotes. Their implication in many cell signaling pathways make them interesting targets for drug development. Three members of this family are nuclear: SIRT1, SIRT6 and SIRT7, where they act as important players for epigenetic regulation through their NAD+ dependent histone deacetylase function^1^. SIRT6, which attracts a lot of interest for its anti-ageing properties^2^, exhibits structural and functional specificities with respect to the other sirtuins. The structure of sirtuin proteins are made of two main domains, a Rossman fold catalytic domain and a zing finger domain, flanked by disordered N-term and C-term extensions^3^ – see Figure1-a. A helix bundle linking the catalytic domain to the zing finger, conserved in the other sirtuins, is absent in SIRT6. Regarding its function, SIRT6 targets specific acetylated lysine sites on histone H3 tails: H3K9ac, H3K18ac, H3K27ac, H3K56ac^4^. Besides, it was observed that SIRT6 poorly deacetylates free histones, and is specifically functional towards nucleosome substrates. Recent cryogenic electron microscopy (cryoEM) investigations revealed that SIRT6 structural particularity might drive this substrate specificity by using the zinc finger domain as a key anchor on the nucleosome H2A-H2B acidic patch, with the lack of connecting helix bundle providing enough flexibility for the catalytic domain to bind to and move with the DNA helix terminus, poising itself adequately for histone H3 deacetylation^5–7^. SIRT6 binding to the nucleosome core particle induces unwrapping of the DNA entry/exit at the binding site. Molecular dynamics simulations have shown that this local unwrapping can favor the accessibility of buried lysine sites located close to the H3 core (e.g., H3K27, H3K18) by proving enough space for the H3 tail to protrude between the open DNA helix and the histone core, positioning these lysines closer to SIRT6 active site^5^. In a very recent work, Cole’s group have reported cryoEM structures of SIRT6 bound to an acetylated NCP that support with our theoretical observations, further validating that DNA unwrapping is indeed necessary for SIRT6 to access H3K27ac but not H3K9ac^8^.

**Figure 1.**
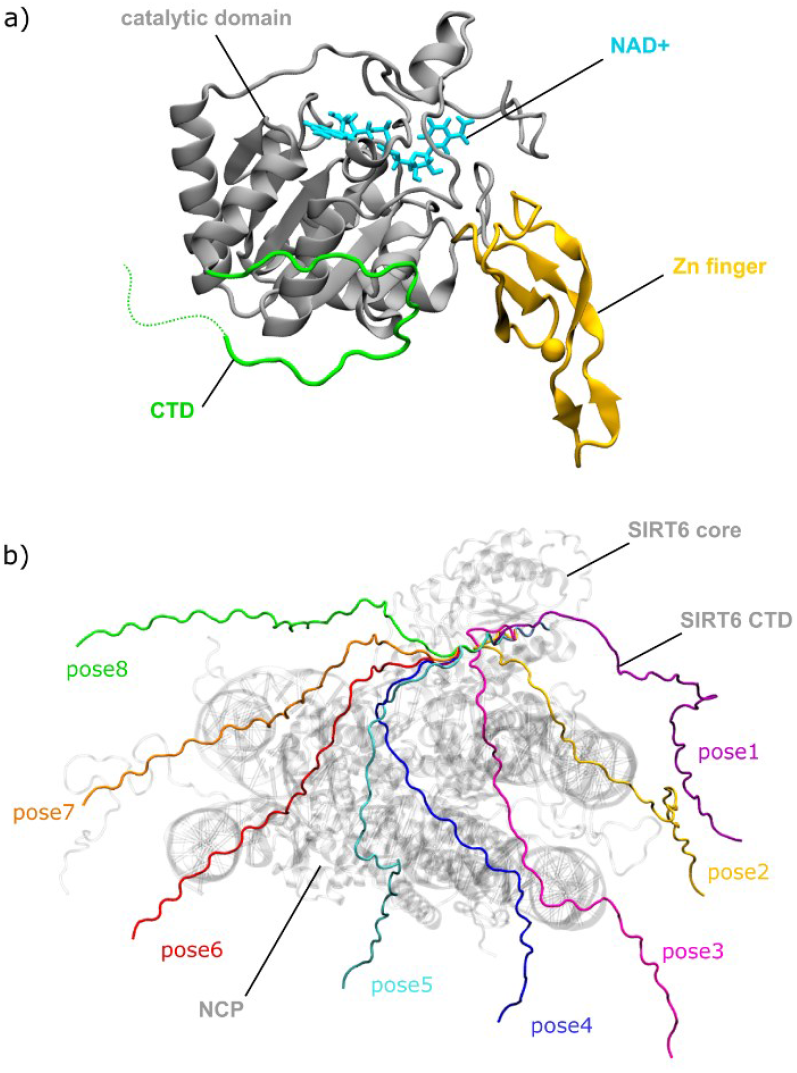
**a)** SIRT6 structure. a catalytic domain structured as a Rossman fold (grey) that harbors the NAD+ cofactor (cyan), a Zinc-finger domain (yellow) that anchors the enzyme to the nucleosome acidic patch, and a C-terminal domain (CTD in green, not fully displayed here). **b)** Top view of the full length SIRT6:NCP complex model, showing the 8 different orientations of the CTD used as starting points for MD simulations.

While these invaluable insights shed light on the characteristics of SIRT6 binding to a nucleosome core particle, they could not provide any structural information about the C-terminal domain of the enzyme. This extended domain is made of 83 amino acids, sharing little sequence identity with other sirtuins, and exhibits a disordered behavior which renders it elusive to structural biology approaches. Experimental investigations of SIRT6 domains function suggested that the CTD could play an important role for the recognition of nucleosomal DNA, high affinity engagement and acetylated lysine processing^9^. However, there is a lack of structural information that would help to understand how SIRT6 CTD actually interacts with DNA in the nucleosome architecture, and how this binding could affect the SIRT6:NCP dynamics and lysine accessibility towards deacetylation.

In this work, we resort to extensive classical MD simulations to explore the binding modes of the SIRT6 CTD to a nucleosome core particle structure, and its effect on DNA unwrapping and lysine accessibility to the enzyme active site. Starting from 8 different orientations of the CTD (see Figure 1-b), we explored its interactions with the DNA helix and the histones, in order to probe how could impact the overall complex dynamics in the context of SIRT6-catalyzed H3 deacetylation. Our results show that SIRT6 CTD tends to favor interactions with the DNA entry/exit on the nucleosome side where the enzyme binds. These binding modes can promote DNA unwrapping and accessibility to lysine sites close to the histone core and barely reachable without DNA opening, suggesting that the CTD might play an important role in H3K27 deacetylation by SIRT6. This contribution brings new insights into the dynamical behavior of the full-length SIRT6 upon binding to a nucleosome core particle. We describe various CTD binding modes to the nucleosome and their effect on lysine accessibility, bringing out new information rationalizing the ability of this enzyme to process multiple lysine sites along the histone H3 tail.

## MATERIALS AND METHODS

### Systems set up

The structure of the full length holo human SIRT6 bound to the NCP was prepared from the cryoEM structure of the apo SIRT6:NCP complex (PDB ID 8OF4^5^). The NAD+ was placed in active site with respect to the ADP ribose moiety coordinates as found in the SIRT6:ADPr complex structure (PDB ID 3PKI^3^). The CTD residues missing in the cryoEM structure (286 to 355) were taken from the Alphafold model of the full-length SIRT6 (AF-Q8N6T7-F1-v4). Given the length of the CTD and its disordered properties, eight different starting structures were generated exhibiting different orientations with respect to the nucleosome, in order to maximize the sampling of its possible conformations in the complex - see Figure 1-b. Protein residues were protonated according to pKa predictions from propKa3^10^. Parameters for the protein and the nucleic acids were taken from the Amber ff14SB^11^ and bsc1force fields^12^, supplemented by CUFIX corrections to improve electrostatics^13^. The CTD disordered region was described with the ff14SBIDPSFF force field^14^ using the CMAP method^15^. Parameters for the zinc finger and the NAD+ were taken from the literature^16,17^. The complex was soaked in a TIP3P water box shaped as a truncated octahedron using a 20Å buffer. 420 Na+ and 287 Cl-ions were added to neutralize the total charge and to ensure a 0.15M ionic concentration, resulting in systems accounting for 670k to 789k atoms.

### Molecular Dynamics simulations and structural analysis

All MD simulations were performed with the Amber20 suite of programs^18^. Starting structures were optimized through a 20,000 steps minimization, and thermalized from 0 to 300K along a 40 ps NVT run using the Langevin thermostat with a 2ps^−1^ collision frequency. The solute atoms were kept restrained using a 10 kcal/mol force constant in these first steps. The systems were then equilibrated during 20 ns in NPT, followed by a final 6 µs production run in NPT. The sampling of one of the replicates featuring the CTD starting orientation far from the DNA entry/exit (pose6) was extended to 11 µs to probe the convergence of the binding mode observed on a longer time scale. The time step was set to 4 fs by taking advantage of the Hydrogen Mass Repartitioning tool^19^. The Particle Mesh Ewald approach was used to treat electrostatic contributions using a 8Å cutoff. 3×5µs control simulations of both the isolated NCP and NCP:SIRT6 complex without the CTD were taken from a previous study^5^, resulting in a total amount 73 µs MD data.

Structural analysis of the MD ensembles was performed using the cpptraj tool implemented in Amber20. The amount of unwrapping was probed using two descriptors reflecting DNA opening with respect to the nucleosome plane: the θ angle connecting SHL0, SHL-5 and SHL-7, and the φ dihedral angle connecting SHL0, SHL2.5, SHL-5 and SHL-7 to quantify the in-plane opening. Distances between the target lysine sites and SIRT6 active sites were measured taking SIRT6:H133 center of mass a reference for the active site, as NAD+ was missing from control simulations without the CTD taken from a previous work^5^. Binding free energies of the SIRT6:NCP complex were estimated using the MMGBSA approach implemented in Amber20. The estimations were performed on 50 frames equally spaced on the last 500 ns of the simulation, for each replicate. Visual inspection and snapshot rendering of the MD trajectories were performed using VMD1.9.3^20^.

## RESULTS AND DISCUSSION

### SIRT6 CTD interacts preferentially at the DNA entry/exit site and can promote unwrapping

In order to assess how SIRT6 CTD interacts with the nucleosome in the NCP:SIRT6 complex, 6 µs MD simulations were performed from 8 different starting orientations – see Figure 1-b. The MD data reveal that the CTD can adopt various binding modes, and that it tends to favor interactions with nucleosomal DNA, in line with previous experimental observations^9^. This behavior reflects the positively charged, disordered properties of this domain. The MD results allow to pinpoint more precisely how the CTD interacts with the nucleosomal DNA. It shows a high propensity of this disordered extension to bind at the DNA entry/exit site near SIRT6 catalytic core docking area, where binding free energies estimations also show the most favorable trends (poses 1-4 and 6) – see Figure 2-a and Table S1.

**Figure 2.**
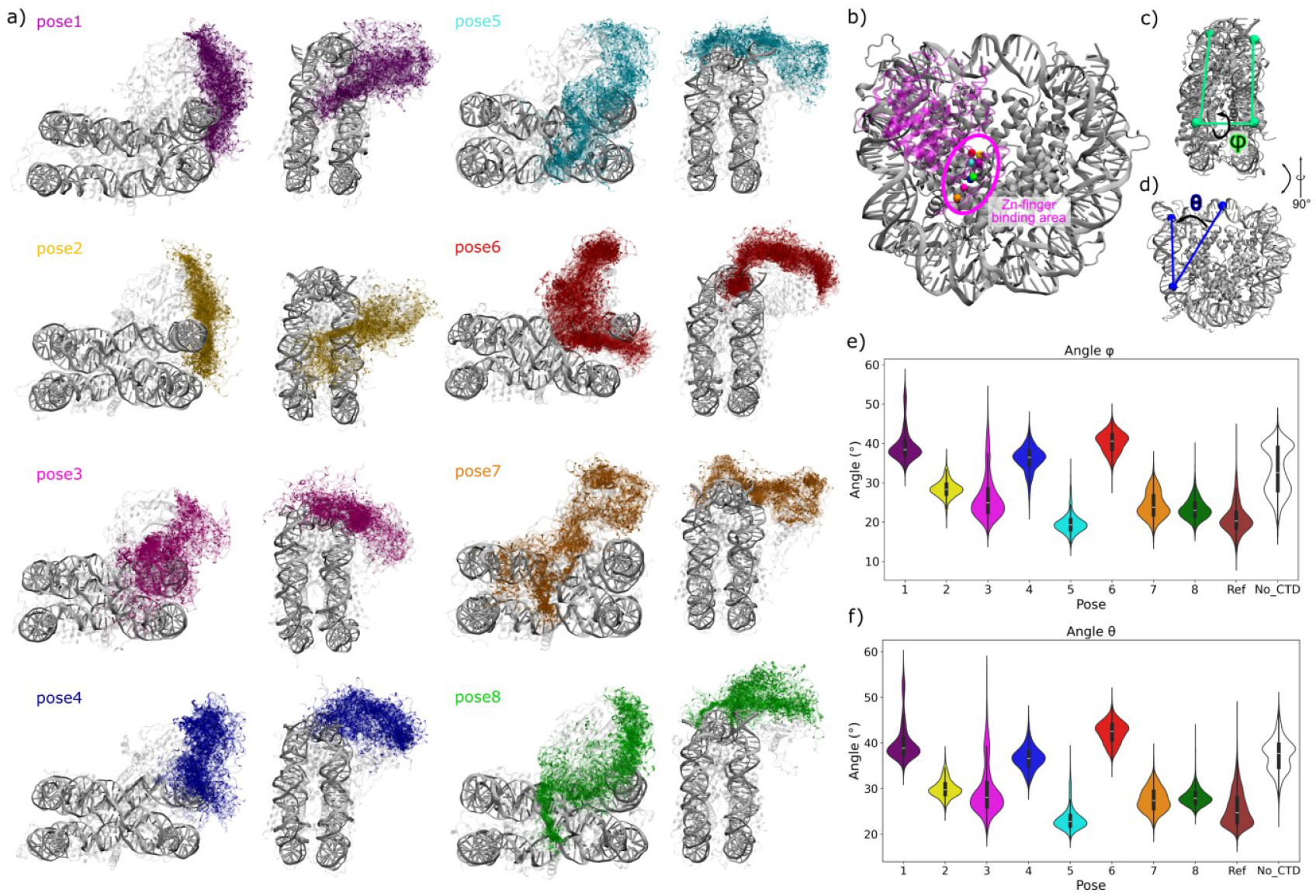
**a)** Distribution of the CTD conformations in the NCP:SIRT6 complex for each starting pose. Conformations are displayed with a 100 ns increment, and the histones and the structured domains of SIRT6 are shown in transparent. **b)** Position of the Zn^2+^ ion in the last frame of the simulation for each replicate (spheres colored according to the starting pose, SIRT6 in transparent magenta). **c)** Definition of the φ dihedral angle and **d)** the θ angle used to quantify the DNA opening. **e)** Distribution of the φ and **f)** θ angles calculated for each replicate (poses 1 to 8), for the isolated NCP (Ref, brown) and for the SIRT6:NCP complex lacking the CTD (No_CTD, white).

The previously-described multivalent interactions of the full-length SIRT6 to the nucleosome were globally found to be stable along the simulations, independently of the CTD starting orientation. The zing finger domain exhibited very slight deviations, but remained anchored to the H2A-H2B acidic patch – see Figure 2-b. The key R220, R231 and R248 residues of the Rossman fold domain were found to consistently make contact with the DNA at SHL6-6.5 as described in a recent cryoEM structure of SIRT6 bound to a H3K27ac-bearing NCP^8^. However, the H2A C-terminal tail showed more fluctuations, switching between interactions with SIRT6 N-terminal domain and its zinc finger domain, but stayed in contact with SIRT6 over the whole MD ensembles. Previous experimental studies suggested various implications for the H2A C-term in the SIRT6:NCP complex: interplay with H2A ubiquitination^5^, inhibitory interactions with SIRT6^6^, and overall stabilization of the complex^7^. The recent cryoEM work by Cole’s lab suggests that the H2A tail is labile and that its interaction with SIRT6 is not required for the enzyme’s function^8^, in good agreement with our observations.

To understand if the CTD could have any role in SIRT6 deacetylase activity other than its contribution to nucleosome binding, DNA unwrapping was measured using the φ and θ angles described above – see Figure 2c-f. Interestingly, we observed that the CTD binding mode drives its effect on DNA unwrapping, with contrasted results obtained for some of the poses. Overall, the DNA opening mostly happens in the plane of the nucleosome wheel (same global trends for φ and θ), probably because of the presence of SIRT6 catalytic domain which hinders out-of-plane movements. As a comparison, DNA unwrapping observed upon redox modification of the histone H3 in isolated NCP simulations showed out-of-plane opening^21^.

The conformations promoting DNA unwrapping, observed in poses 1, 4 and 6, exhibit direct interaction with the DNA extremity, yet adopting various binding modes – see Figures 2 and S1. In pose 1, the 15 last residues of the CTD, harboring 3 lysines (K346, K349, K351) and 2 arginines (R343, R347), form very stable interactions in the DNA minor groove at SHL-6. The largest opening angles are observed for this binding mode ( φ_max_= 57° and θ_max_= 59°, see Figure 2-e and f), also favoring the protrusion of the H3 tail between the DNA end and the histone core – see details in section below. The upstream part of the CTD interacts transiently with the DNA major groove, next to the H3 tail end. The CTD can also interact with the extremity of the DNA helix, as seen in pose 4, and favor DNA unwrapping as illustrated by characteristic φ and θ values for open DNA conformations. In the 11µs-long replicate corresponding to pose 6, the CTD initially points towards the DNA end opposite to SIRT6 binding site. Interestingly, it moves rapidly towards the other DNA end, i.e. near SIRT6 core, and interact with the DNA helix in yet a different way than for poses 1 and 4, by placing itself in between the DNA helix and the histone H3 core. This binding mode exhibits the highest average values of φ and θ, probably by directly destabilizing the DNA-histone core interactions. Noteworthy, in these three situations, the basic residues that make most of the interactions with the DNA are the ones located at the terminus of the CTD, suggesting that they might be crucial for CTD-mediated SIRT6-NCP interactions and the overall stability of the complex.

For some other binding modes, i.e. poses 2 and 3, the CTD have no effect on DNA unwrapping with respect to the control simulations lacking this domain. In both cases, the basic residues of the CTD end do not form any stable interaction with the DNA helix. In pose 2, the CTD binds in the minor groove at SHL-6 via K332, R333 and R335 located slightly upstream of the terminal basic residues on the CTD – see Figure S1. While these interactions are stable, they hinder DNA unwrapping. The involvement of these residues instead of the non-terminal basic ones might act as a more constrained latch, hampering large movements of the DNA helix. For pose 3, the CTD moves towards SIRT6 core but only transiently interacts with the DNA helix. While φ and θ can briefly peak at high values during the simulation, their average trends remain comparable to what it observed for the control system lacking CTD.

Finally, some CTD conformations can preclude DNA unwrapping, exhibiting φ and θ values similar to what is observed for the control isolated NCP, even lower than for the SIRT6:NCP complex without CTD. These correspond to binding modes where the CTD does not interact with the DNA entry/exit but rather with DNA at the dyad (poses 5, 7 and 8). Importantly, the free energies of binding estimated for these interaction modes are much lower than for the other poses, suggesting less favorable interactions for the complex stability – see Table S1.

Overall, these results suggest that SIRT6 CTD can adopt a variety of different binding modes, which induce contrasted effects on DNA unwrapping. This domain tends to favor interactions with the DNA end near SIRT6 docking site which, when mediated by the terminal basic residues (R343, K346, R347, K349 and K352), can promote the opening of the DNA end.

### SIRT6 CTD can facilitate the accessibility to H3K27

In a previous study, we showed that the binding of SIRT6 without CTD to the NCP could favor DNA unwrapping, and the protrusion of the H3 tail in between the DNA gyre and the histone core, facilitating the access to buried target lysine sites^5^. After describing how specific binding modes of the CTD in the full length SIRT6:NCP complex could promote the unwrapped states of nucleosomal DNA, we further probed how this could also affect the accessibility of the H3 lysines targeted by the deacetylase.

Overall, the presence of the CTD does not favor the accessibility of the target lysine sites (H3K9, H3K18, H3K27, H3K56) more than in the control complex lacking this domain – see Figure 3-a. The distribution of the distances between each lysine and the catalytic SIRT6:H133 residue show that the CTD has very little effect on the accessibility of the target lysines to its active site. This was not surprising as it has been showed experimentally that SIRT6 deacetylase function does not require the presence of its CTD^9^. It is important to underline that the present simulations do not include any acetylation site, which might influence the observed trends.

**Figure 3.**
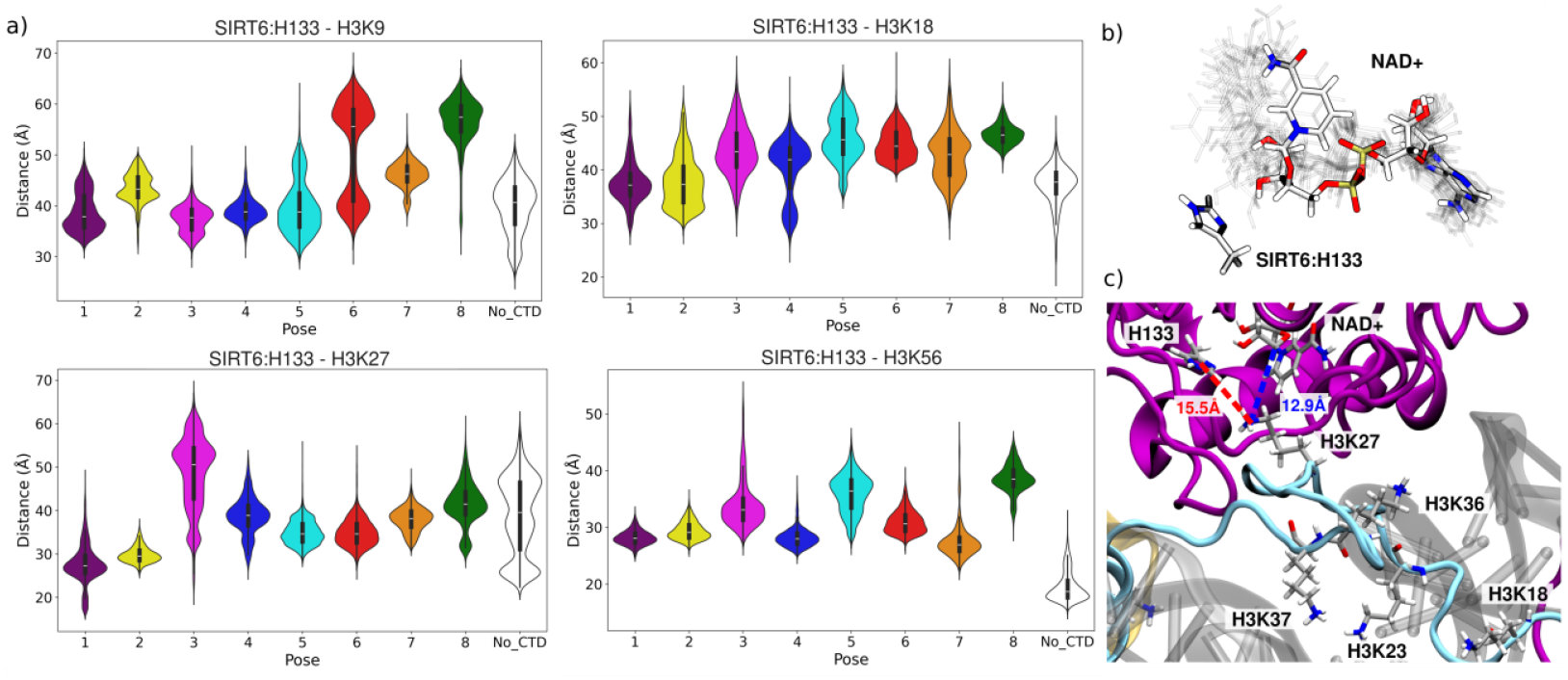
**a)** Distribution of the distance between the SIRT6:H133 catalytic residue and the four lysines processed by the enzyme, for each starting pose (color code corresponding to Figure 1) and for the SIRT6:NCP complex in the absence of CTD (No_CTD, white). **b)** Dynamics of the NAD+ within the active site in pose 6. Twelve conformations are showed in ghost representation that span frames from 1 ns to 5500 ns, and the last frame (t=6000 ns) is showed in colored licorice. **c)** Conformation exhibiting the lowest SIRT6:H133-H3K27 distance observed in the MD from pose 6. The position of the other H3 lysine are also showed, and the H3 and SIRT6 structures are displayed in light blue and magenta, respectively.

An exception is observed for H3K27 in the pose 1 simulation. As described above, this replicate exhibits the highest values of the φ and θ opening angles. In this replicate, the NAD+ cofactor exhibits a strong positioning of its adenosine moiety (see Figure 3-b), ensured by strong interactions with the surrounding T57, N240 and L241 residues. However, the nicotinamide extremity shows a labile behavior, yet staying close to the catalytic H133. Such flexibility might be of importance to accommodate the different lysine sites and post-translational modification types within the active site for processing. The promotion of the DNA unwrapping strongly favors the protrusion of the H3 tail between the DNA helix and the histone core, allowing H3K27 to eventually move closer to SIRT6 active site (minimal distance d_min_= 15.5 Å for H133-H3K27 in pose 1 vs 21.2 Å in the control). Interestingly, this conformation exhibits structural determinants that could resemble a pre-reactive state of the active site – see Figure 3-c. H3K27@NZ is positioned 12.9 Å away from the reactive NAD+@C15 atom, while the nitrogen atom of SIRT6:H133 that captures a proton from NAD+ during the reaction is located only 3.1 Å away from it. This conformation constitutes an excellent starting point for further *in silico* studies of the SIRT6:NCP complex harboring H3K27ac, that would shed light on the mechanisms driving the processing of this buried site by the deacetylase. The other target sites (H3K9, H3K18, H3K56) do not spontaneously come close to SIRT6 active site, and might necessitate more complex dynamical rearrangements to be processed for deacetylation.

## CONCLUSIONS

The dynamical behavior of the C-terminal domain of the SIRT6 deacetylase was investigated to understand its role in nucleosome binding and histone deacetylation. Exploiting a total of 73 µs MD simulations data, we reveal a variety of binding modes of the CTD onto nucleosomal DNA, exhibiting contrasted effects on DNA unwrapping and lysine accessibility. We highlight the high propensity of the CTD to interact with the DNA entry/exit near the SIRT6 core docking site, through conformations associated with the most favorable binding free energy estimations compared to other binding modes. Our results suggest that highly basic terminal residues on the CTD end (i.e. R343, K346, R347, K349, K351) interaction with the DNA end is of major importance for the stabilization of unwrapped DNA states. The promotion of DNA unwrapping upon the presence of the CTD allowed the appearance of specific conformations featuring H3K27 near the active site (<13Å), in agreement with recent cryoEM structure of SIRT6:NCP in the presence of H3K27ac^8^. This work provides an important starting point for combined MM- and QM/MM-MD studies to further shed light on SIRT6 fine-tuned specificity, and offers invaluable structural insights into the mechanisms driving H3K27 deacetylation by SIRT6.

## ASSOCIATED CONTENT

## Supporting Information

The following files are available free of charge:

Supplementary figure and table (PDF)

MD trajectories files are available on Zenodo at https://doi.org/10.5281/zenodo.15010369

## Author Contributions

The manuscript was written through contributions of all authors. All authors have given approval to the final version of the manuscript.

## ACKNOWLEDGMENT

EB thanks the IDRIS and CINES centers for computational resources (allocation 2023-A0150714577) made by GENCI.

## ABBREVIATIONS

CTD: C-Terminal Domain
CryoEM: Cryogenic Electron Microscopy
MD: Molecular Dynamics
NCP: Nucleosome Core Particle
SIRT6: Sirtuin 6

## TABLE OF CONTENTS GRAPHIC

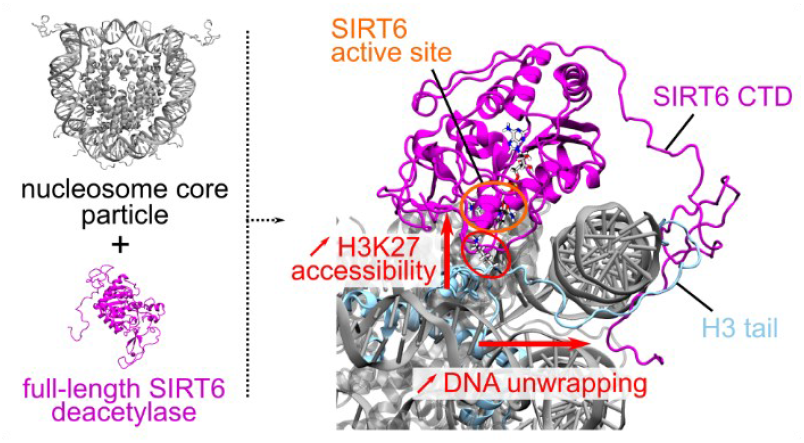

## Notes

https://doi.org/10.5281/zenodo.15010369

### Competing Interest Statement

The authors have declared no competing interest.

## REFERENCES

(1) Wątroba, M.; Dudek, I.; Skoda, M.; Stangret, A.; Rzodkiewicz, P.; Szukiewicz, D. Sirtuins, Epigenetics and Longevity. Ageing Research Reviews 2017, 40, 11–19. 10.1016/j.arr.2017.08.001.

(2) Kanfi, Y.; Naiman, S.; Amir, G.; Peshti, V.; Zinman, G.; Nahum, L.; Bar-Joseph, Z.; Cohen, H. Y. The Sirtuin SIRT6 Regulates Lifespan in Male Mice. Nature 2012, 483 (7388), 218–221. 10.1038/nature10815.

(3) Pan, P. W.; Feldman, J. L.; Devries, M. K.; Dong, A.; Edwards, A. M.; Denu, J. M. Structure and Biochemical Functions of SIRT6. J Biol Chem 2011, 286 (16), 14575–14587. 10.1074/jbc.M111.218990.

(4) Wang, W. W.; Zeng, Y.; Wu, B.; Deiters, A.; Liu, W. R. A Chemical Biology Approach to Reveal Sirt6-Targeted Histone H3 Sites in Nucleosomes. ACS Chem. Biol. 2016, 11 (7), 1973–1981. 10.1021/acschembio.6b00243.

(5) Smirnova, E.; Bignon, E.; Schultz, P.; Papai, G.; Ben-Shem, A. Binding to Nucleosome Poises Human SIRT6 for Histone H3 Deacetylation. bioRxiv July 18, 2023, p 2023.03.20.533444. 10.1101/2023.03.20.533444.

(6) Chio, U. S.; Rechiche, O.; Bryll, A. R.; Zhu, J.; Leith, E. M.; Feldman, J. L.; Peterson, C. L.; Tan, S.; Armache, J.-P. Cryo-EM Structure of the Human Sirtuin 6-Nucleosome Complex. Sci Adv 2023, 9 (15), eadf7586. 10.1126/sciadv.adf7586.

(7) Wang, Z. A.; Markert, J. W.; Whedon, S. D.; Yapa Abeywardana, M.; Lee, K.; Jiang, H.; Suarez, C.; Lin, H.; Farnung, L.; Cole, P. A. Structural Basis of Sirtuin 6-Catalyzed Nucleosome Deacetylation. J Am Chem Soc 2023, 145 (12), 6811–6822. 10.1021/jacs.2c13512.

(8) Wang, Z. A.; Markert, J.; Whedon, S. D.; Abeywardana, M. Y.; Sheng, X.; Nam, E.; Lee, K.; Chen, M.; Waterbury, A.; Zhao, Y.; Farnung, L.; Cole, P. A. Structural and Enzymatic Plasticity of SIRT6 Deacylase Activity. Journal of Biological Chemistry 2025, 108446. 10.1016/j.jbc.2025.108446.

(9) Liu, W. H.; Zheng, J.; Feldman, J. L.; Klein, M. A.; Kuznetsov, V. I.; Peterson, C. L.; Griffin, P. R.; Denu, J. M. Multivalent Interactions Drive Nucleosome Binding and Efficient Chromatin Deacetylation by SIRT6. Nat Commun 2020, 11 (1), 5244. 10.1038/s41467-020-19018-y.

(10) Olsson, M. H. M.; Søndergaard, C. R.; Rostkowski, M.; Jensen, J. H. PROPKA3: Consistent Treatment of Internal and Surface Residues in Empirical p K _a_ Predictions. J. Chem. Theory Comput. 2011, 7 (2), 525–537. 10.1021/ct100578z.

(11) Maier, J. A.; Martinez, C.; Kasavajhala, K.; Wickstrom, L.; Hauser, K. E.; Simmerling, C. ff14SB: Improving the Accuracy of Protein Side Chain and Backbone Parameters from ff99SB. Journal of chemical theory and computation 2015, 11 (8), 3696–3713.

(12) Ivani, I.; Dans, P. D.; Noy, A.; Pérez, A.; Faustino, I.; Hospital, A.; Walther, J.; Andrio, P.; Goñi, R.; Balaceanu, A.; Portella, G.; Battistini, F.; Gelpí, J. L.; González, C.; Vendruscolo, M.; Laughton, C. A.; Harris, S. A.; Case, D. A.; Orozco, M. Parmbsc1: A Refined Force Field for DNA Simulations. Nat Methods 2016, 13 (1), 55–58. 10.1038/nmeth.3658.

(13) Yoo, J.; Aksimentiev, A. New Tricks for Old Dogs: Improving the Accuracy of Biomolecular Force Fields by Pair-Specific Corrections to Non-Bonded Interactions. Phys. Chem. Chem. Phys. 2018, 20 (13), 8432–8449. 10.1039/C7CP08185E.

(14) Song, D.; Luo, R.; Chen, H.-F. The IDP-Specific Force Field ff14IDPSFF Improves the Conformer Sampling of Intrinsically Disordered Proteins. J Chem Inf Model 2017, 57 (5), 1166–1178. 10.1021/acs.jcim.7b00135.

(15) MacKerell, A. D.; Feig, M.; Brooks, C. L. Improved Treatment of the Protein Backbone in Empirical Force Fields. J Am Chem Soc 2004, 126 (3), 698–699. 10.1021/ja036959e.

(16) Macchiagodena, M.; Pagliai, M.; Andreini, C.; Rosato, A.; Procacci, P. Upgraded AMBER Force Field for Zinc-Binding Residues and Ligands for Predicting Structural Properties and Binding Affinities in Zinc-Proteins. ACS Omega 2020, 5 (25), 15301–15310. 10.1021/acsomega.0c01337.

(17) Walker, R. C.; de Souza, M. M.; Mercer, I. P.; Gould, I. R.; Klug, D. R. Large and Fast Relaxations inside a Protein: Calculation and Measurement of Reorganization Energies in Alcohol Dehydrogenase. J. Phys. Chem. B 2002, 106 (44), 11658–11665. 10.1021/jp0261814.

(18) Case, D.; Ben-Shalom, I.; Brozell, S.; Cerutti, D.; Cheatham III, T.; Cruzeiro, V.; Darden, T.; Duke, R.; Ghoreishi, D.; Gilson, M.; others. AMBER 2018: San Francisco, 2018.

(19) Hopkins, C. W.; Le Grand, S.; Walker, R. C.; Roitberg, A. E. Long-Time-Step Molecular Dynamics through Hydrogen Mass Repartitioning. Journal of chemical theory and computation 2015, 11 (4), 1864–1874.

(20) Humphrey, W.; Dalke, A.; Schulten, K. VMD: Visual Molecular Dynamics. Journal of molecular graphics 1996, 14 (1), 33–38.

(21) Karami, Y.; González-Alemán, R.; Duch, M.; Qiu, Y.; Kedjar, Y.; Bignon, E. Histone H3 as a Redox Switch in the Nucleosome Core Particle: Insights from Molecular Modeling†. October 7, 2024. 10.1101/2024.10.07.616940.

