## Supplementary table and figure for "Specific binding modes of SIRT6 C-terminal domain to the nucleosome core particle influence DNA unwrapping and H3K27 accessibility"

**Table S1.** MM/GBSA binding free energy estimations of the full-length SIRT6:NCP complex for each pose, calculated on the last 500 ns of the simulations.

| Replicate | Average $\Delta G_{binding}$ (kcal/mol) | Standard deviation (kcal/mol) |
|---|---|---|
| Pose 1 | -135.7 | 19.0 |
| Pose 2 | -132.2 | 17.8 |
| Pose 3 | -140.0 | 14.6 |
| Pose 4 | -90.1 | 13.9 |
| Pose 5 | -69.2 | 13.2 |
| Pose 6 | -92.4 | 17.0 |
| Pose 7 | -60.1 | 18.3 |
| Pose 8 | -53.1 | 14.6 |

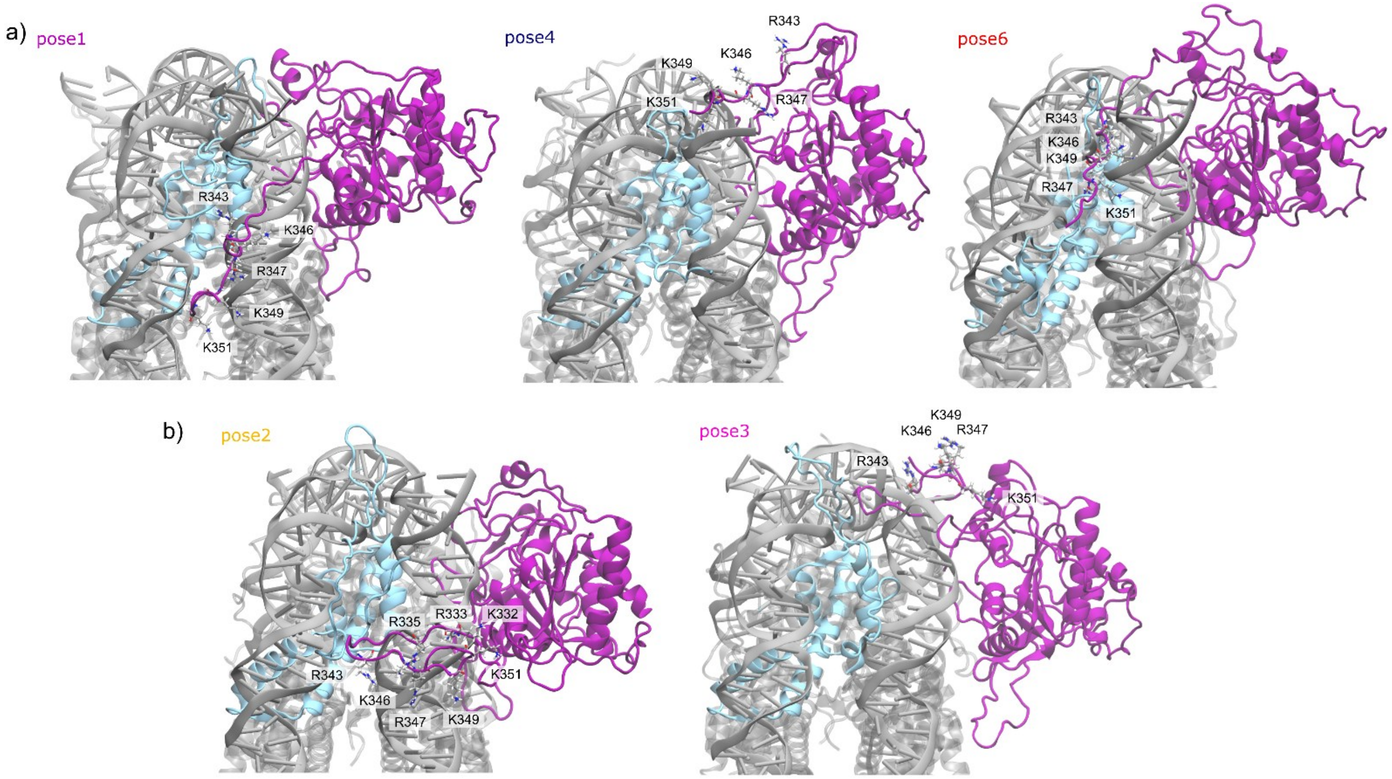


**Figure S1.** Side view of the SIRT6:NCP complex, showing the conformation of the CTD end (last frame) for **a)** binding modes favoring DNA unwrapping (poses 1, 4, and 6) and **b)** binding modes that do not influence DNA unwrapping (poses 2 and 3). The basic residues located at the end of the CTD (R343, K346, R347, K349 and K352) are displayed in licorice.
